# Imputed transcriptional patterns in ancient genomes reveal molecular targets of selection

**DOI:** 10.1101/2025.11.21.689732

**Authors:** Mathilde André, Francesco Montinaro, Ettore Zapparoli, Paolo Provero, Davide Marnetto

**Affiliations:** Dept. of Neurosciences “Rita Levi Montalcini”, University of Turin, Torino, Italy; Dept. of Biosciences, Biotechnology and Environment, University of Bari, Bari, Italy; Institute of Genomics, University of Tartu, Tartu, Estonia

**Author notes:** Correspondence to: Mathilde André (.) and Davide Marnetto.

**Keywords:** Genetic expression, splicing, ancient DNA, natural selection

## Abstract

The genetically regulated component of molecular traits can be imputed from ancient genomes, though low genotyping quality and genetic drift may impair accurate expression prediction and results interpretation. We developed tissue-specific gene expression and alternative splicing prediction models and applied them to over a thousand imputed ancient genomes spanning 10,000 years of Western Eurasian history. We revealed gene regulatory patterns diverging with time, space and genomic ancestry by fitting predicted transcriptional profiles onto multivariable models including these factors. By interpreting associations with time as recent natural selection we identify hits mediated by transcriptional changes. Finally, we fine-mapped these selection signals to tissue-specific transcriptional traits, explicitly accounting for prediction uncertainty. Among others, we elucidate the spatial patterns and the regulatory features promoting selection at *LCT, FADS1, SLC22A5/*P4HA2, *ACAD10*/*OAS3* and find novel signals of selection at *TBIMB6, SORD2P* and *AC012370*.*3*.

## Introduction

Present-day Western Eurasian populations reflect a complex history of migration and admixture^1–4^. Over the last decade, the increasing availability of ancient genomic datasets from this region has been essential in understanding population dynamics^1–6^ and how major shifts in lifestyle and environment have shaped both ancient and modern Eurasian genomes through natural selection^7–10^. However, identifying the target variants under selection and the underlying molecular mechanism or phenotype is often challenging, hindering a deep understanding of our evolutionary past.

A substantial fraction of selection signatures reside in non-coding regions of the genome and are likely associated with gene regulatory functions^11^. Consequently, the annotation of selection candidates through molecular quantitative trait loci^12,13^, especially gene expression and splicing, offers a promising approach to infer selection-driving mechanisms^11,14,15^.

As learning methods that leverage large-scale training data can predict molecular phenotypes from genetic variation^16–18^, imputing gene expression from ancient genomes and tracking it through time and populations emerged as an alternative strategy^19^. For example, Colbran et al.^20^ explored whether predicted expression levels could be associated with lifestyle transitions, while Poyraz et al.^21^ examined temporal dynamics of expression in ancient Britain. These two studies provided foundational insights into the evolution of gene regulation in Europe, but had to tackle the relatively low genotyping quality of ancient DNA (aDNA), which limits genotyping resolution and restricts sampling. In particular, Colbran et al.^20^ were constrained by the coverage of the 1240k variants assayed in ancient humans from the Allen Ancient DNA Resource (AADR)^22^. Poyraz et al.^21^, albeit employing an advanced imputation and expression prediction pipeline to overcome limitations of the previous study, focused on a narrow spatial and temporal interval: the last 4,500 years in Britain.

In this work, we expand upon these earlier efforts along three axes. First, we leverage a newly released imputed ancient genome panel spanning the last 10,000 years across Eurasia^6^, increasing both the genotyping density and spatio-temporal range of sampled individuals. Second, we include alternative splicing predictions, adding a previously unexplored regulatory dimension. Third, by explicitly addressing the relationship across transcriptional features in different tissues, we aimed to finemap the selection signal onto a specific tissue/transcript combination. Collectively, these advances allow us to revisit known loci under selection, identify new ones, and provide novel insight into the transcriptomic basis of adaptation in ancient western Eurasian populations.

## Results

### Imputing transcriptional patterns in ancient samples

We predicted genetically regulated gene expression (GReX) and alternative splicing (GReS) across 14 tissues in 1,145 imputed ancient Western Eurasian genomes^6^ plus 80 modern European samples from the 1000 Genomes panel^23,24^, later referred to as the “discovery set” (**Figure S1, Table S1**). GReX and GReS (GRe*) predictions were obtained using an elastic net regression model^16,17^ trained on samples of European ancestry from GTEx^13^. After removing models with low predictivity in a hold-out GTEx set (R < 0.1) or poor imputation at predictive variants in the discovery set, we predicted a total of 252,231 GRe*, with an average of 6,065 GReX and 11,952 GReS per tissue (**Table S2, S3**).

In the first of the two studies introduced above, Colbran et al.^20^ downsampled the GTEx training set to the 1240k variants assayed in ancient humans from the Allen Ancient DNA Resource (AADR)^22^ to adapt their prediction models to ancient samples, . Because a fully imputed dataset of ancient genomes is now available, we were able to use full GReX models: indeed 95.02% of the GReX-predictive SNPs (SNP coefficient != 0), which were reliably imputed (mean info score = 0.878), are available in our discovery set in comparison with 17.46% found in the 1240k capture array. Nonetheless, when downsampling our training set as suggested in Colbran et al.^20^ and comparing with the full model, we observed a highly correlated performance for GTEx European (Pearson’s ρ = 0.95) and Non-European samples (Pearson’s ρ = 0.89). Despite this strong correlation we observed a significant improvement of the performance within both groups (P = 3.99e-74 and P = 1.15e-43, respectively; two-tailed paired Wilcoxon test; **Figure S2A**,**B**). In particular, GReX full models tend to perform better for genes for which there is a larger increase in the number of predictive SNPs between the 1240k and the full model (P = 4.59e-25 and P = 7.60e-05, respectively; two-tailed Mann-Whitney U test; **Figure S2C**,**D**). Sun-exposed skin was used as example tissue in this analysis.

### Transcriptome-wide scan for non-neutral changes in time

To identify non-neutral transcriptional patterns across time and space, we regressed each GRe* (i.e. each tissue and gene expression/splicing pair) onto average dating (hereafter time), latitude and longitude, including their pairwise interaction terms, and the first 10 genomic principal components (PCs, **Figure S3**) of each sample. Notably, with such a diverse dataset, PCs are necessary to reduce neutral patterns solely explainable by drift. Indeed, just regressing e.g. the sun-exposed skin GReX of all genes onto time would result in a Genomic Inflation Factor λ=6.64. On the other hand, as recognized by others^25^, PCs are heavily correlated with time (the first 10 PCs explain 80% of the variance of time) so, while ensuring specificity, they presumably hinder the sensitivity of our approach.

A significant association with time in the multivariable regression model, indicating a GRe* directional shift in the time transect analyzed, is interpreted as selection (time coefficients and P-values for all models are reported in **Supplementary Data 1**). Reassuringly, time coefficient P-values are not correlated with either model predictivity (R) or imputation quality at predictive variants in nearly any tissue (**Table S4**). P-values across tissues were combined with ACAT^26^ to have one signal per gene. Resulting P-values for association with time showed an inflation of λ=1.58 and λ=1.89 for GReX and GReS, respectively, so we further corrected them by genomic control (GC). As shown in **Figure 1A** and **1B**, we identified 8 genomic loci each where GReX or GReS are significantly associated with time at 5% False Discovery Rate (FDR), 11 when pooling both GRe*. Of these, 6 loci for each transcriptional dimension overlap with one of 18 known and replicable signals of selection collected from literature^7,8,15,21,27,28^(**Table S5**) while 4 genes are novel findings: *DUOX2* (P=5.59e-05) and *AC012370*.*3* (P=1.43e-4) for GReX, *TMBIM6* (P=7.01e-05) and *DNAJB12* (P=2.07e-4) for GReS. Reassuringly, a known adaptation traceable to a change in SLC45A2^29^ amino acidic sequence does not show any significance in our analysis.

**Figure 1:**
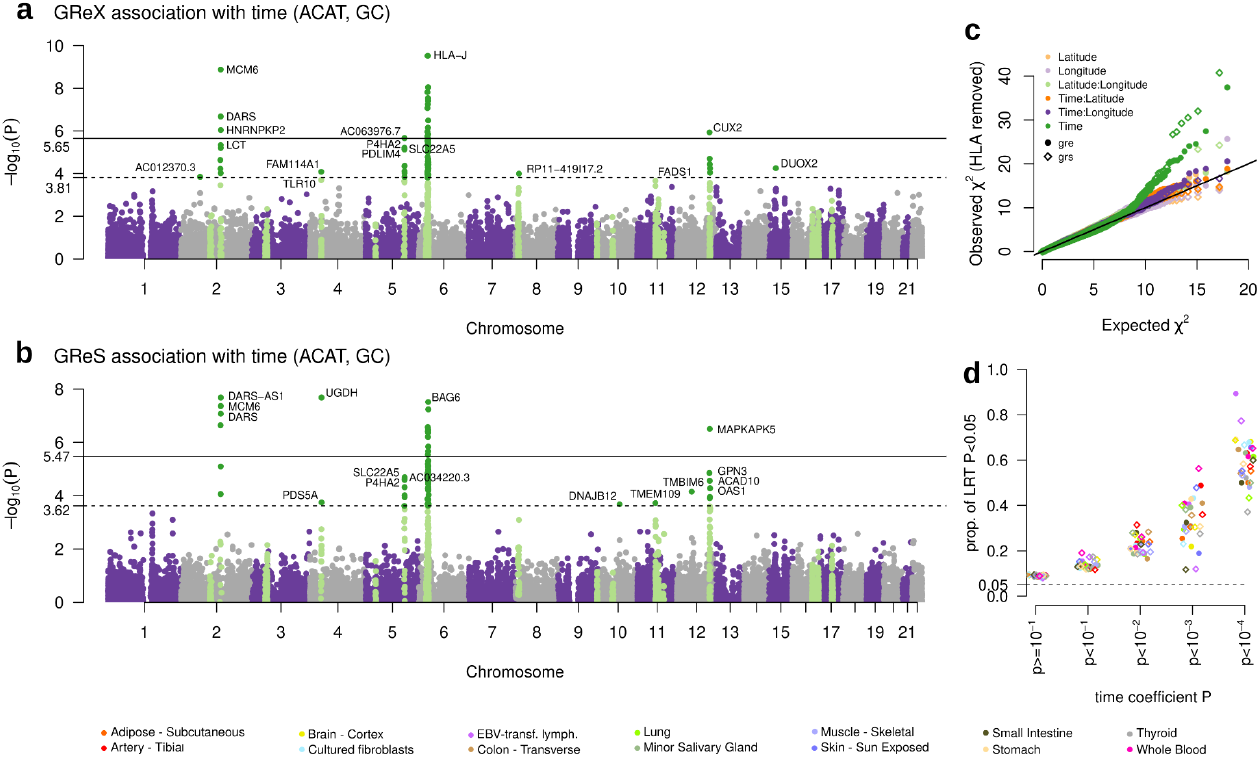
Transcriptome-wide associations with time. **a**. Transcriptome-wide ACAT P-values for association between GReX and time, after genomic control adjustment. A total of 22,277 genes are represented. Genes in light green lie within 18 known and replicable signals of selection collected from literature, while those in dark green are significantly associated with time in our analysis. The solid and dashed lines indicate Bonferroni and 5% FDR significance, respectively **b**. Transcriptome-wide ACAT P-values for association between GReS and time, after genomic control adjustment. 14,868 genes are represented. **c**. QQ plot of *Χ*^2^ for each predictor separately for GReX (dots) and GReS (diamonds) after removing HLA. For this plot including HLA see **Figure S5. d** Proportion of GRe* for each tissue better explained in our discovery dataset by a model including time, latitude, longitude, pairwise interactions and 10 PCs, compared with a model only including time and PCs. GRe* are reported in different columns according to their P-value for association with time. Likelihood Ratio Test, significance at P=0.05, the dashed line represents the 0.05 expectation. Again GReX are dots and GReS diamonds, colors represent tissues.

Associations with time are overall more significant than associations with space and space-time interactions, as visible in the Quantile-Quantile plot in **Figure 1C**; Manhattan plots for all predictors are shown in **Figure S4**. An exception is represented by the HLA locus, containing many GRe* associated with latitude, longitude and their interaction (**Figure S5**). While this locus is surely interesting, its high polymorphism, paralogy, and low recombination makes mapping difficult, hindering the reliability of our approach. We, therefore, chose not to analyze and interpret it further.

Although few signals associated with latitude and longitude reached significance at genome-wide level, these predictors determine an overall better fit of the GRe* predictive models, especially for putatively selected genes (**Figure 1D**) and are instrumental to interpret selection patterns across such a wide spatial range.

### Refinement and fine-mapping of selected loci

By generating tissue-specific GReX and GReS we are able to fine-map selection signals to specific transcriptional features and conditions, disentangling the correlation across GRe*. For 10 significant loci from the previous section (**Table 1**) - all except HLA - we selected all GRe* analyzed before (thus discarding GRe* models with low predictivity and variant imputation quality) and used summary-based SuSiE^30^ with the summary statistics of the GRe* association with time. SuSiE is usually implemented to finemap GWAS trait-variant associations; however, GRe* have greater uncertainty than variant genotypes, which should be taken into account, as underlined by Strober et al.^31^. This is especially important as such uncertainty is feature-dependent, due to differences in GRe* predictability and sample sizes across tissues. We therefore re-trained 10 times every GRe* model on GTEx, leaving out every time a different 20% of the GTEx samples.

**Table 1:** significant loci. Combined effect sizes, P- and Q-values, together with proportion of presence in SuSiE credible sets reported for the most likely candidates identified at each genome-wide significant locus. The GRe* appearing in the largest proportion of SuSiE credible sets and with a significant Q-value are shown for each locus. In case of multiple GRe* summarized in a single row, coefficient P-value and Q-value for the top GRe* are reported. *start of top ACAT gene, hg38; **\*\*** coefficients have been reversed so that positive indicates increasing expression, and negative decreasing. Abbreviations: EJ: exon junction, In: Small intestine, Sk: Sun Exposed Skin, Ms: Skeletal Muscle, Ad: Adipose Tissue, Ar: Tibial Artery, Th: Thyroid.

| locus |  | ACAT |  |  | Refinement |  |  |  |  |
| --- | --- | --- | --- | --- | --- | --- | --- | --- | --- |
| tag | pos (hg38)* | Top gene | Top trait | Top P-value (GC) | Candidate GRe* | Candidate tissue | Effect size avg.** | combined BH Q-value | In CS prop. |
| novel | chr2:65439888 | *AC012370.3* | GReX | 1.43E-04 | *AC012370.3*, expression<br>*AC074391.1*, expression | Thyroid<br>Thyroid | -0.39<br>-0.36 | 8.86E-04<br>3.95E-03 | 0.90<br>0.90 |
| *LCT* | chr2:135839626 | *MCM6* | GReX | 1.33E-09 | *MCM6*, expression | Whole Blood | -0.50 | 7.65E-11 | 0.60 |
| *TLR* | chr4:39498755 | *UGDH* | GReS | 2.06E-08 | *TLR10*, EJ, chr4:38,735,565-38,782,921 | EBV-tr. lymphocytes | 0.31 | 1.64E-02 | 0.20 |
| *IRF1* | chr5:132199456 | *AC063976.7* | GReX | 2.17E-06 | *SLC22A5*, EJ, chr5:132,391,374-132,392,433<br>*P4HA2*, exon, chr5:132,227,086-132,227,221<br>*P4HA2*, expression | Sk, Ms<br>Sk, In, Ms, Ad, Ar, Lung, Th<br>Tibial Artery | -0.48<br>-0.45<br>0.45 | 1.21E-06<br>3.91E-06<br>6.62E-06 | 0.80<br>0.80<br>0.80 |
| HLA | chr6:30005971 | *HLA-J* | GReX | 3.00E-10 | NA | NA | NA | NA | NA |
| *GATA4* | chr8:12476462 | *RP11-419I17.2* | GReX | 1.02E-04 | *RP11-419I17.2*, expression | Transverse Colon | -0.37 | 1.07E-03 | NA |
| novel | chr10:72332830 | *DNAJB12* | GReS | 2.08E-04 | *DNAJB12*, EJ, chr10:72,334,617-72,335,786 | Small Intestine | 0.31 | 2.25E-02 | 0.30 |
| *FADS1* | chr11:60913874 | *TMEM109* | GReS | 1.88E-04 | *FADS1*, expression<br>*FEN1*, expression | Brain Cort.,Stomach,Ad,Th<br>Adipose Subcutaneus | 0.34<br>0.33 | 1.08E-03<br>2.78E-03 | 0.40<br>0.40 |
| novel | chr12:49707725 | *TMBIM6* | GReS | 7.04E-05 | *TMBIM6*, EJ, chr12:49,749,734-49,752,464 | Small Intestine | -0.45 | 1.76E-05 | 1.00 |
| *OAS3* | chr12:111841978 | *MAPKAPK5* | GReS | 3.10E-07 | *ACAD10*, EJ, chr12:111,712,657-111,715,821 | Transverse Colon | 0.53 | 2.38E-08 | 0.63 |
| novel | chr15:45092650 | *DUOX2* | GReX | 5.59E-05 | *SORD2P*, expression | Stomach | -0.40 | 1.77E-04 | 0.77 |

Runs for the same GRe* were combined emphasizing consistency (average Z score, see Methods) leading to more reliable associations for each feature, followed by Benjamini-Hochberg transformation into Q-values considering all 252,231 GRe* tested. While different runs led to broadly consistent associations with time, as visible in **Figure S6A**, using a representative significance threshold of absolute Z>3.89 (corresponding to P<10^-4^ without GC and Q=0.032 in our analysis) 41.3% of GRe* with a significant combined Z would be rejected in at least one run (**Figure S6B**), reiterating the importance of accounting for GRe* uncertainty.

We counted how many times each GRe* appeared in a credible set (CS, a set of features containing the causal one with 95% probability) across 30 SuSiE runs, varying the number of allowed CS from 1 to 3 for each of the 10 GRe* training runs. We report the most likely candidate GRe* for each locus analyzed in **Table 1** and discuss examples in **Figure 2** in the next section (full results in **Table S6** and **Figure S7**).

**Figure 2:**
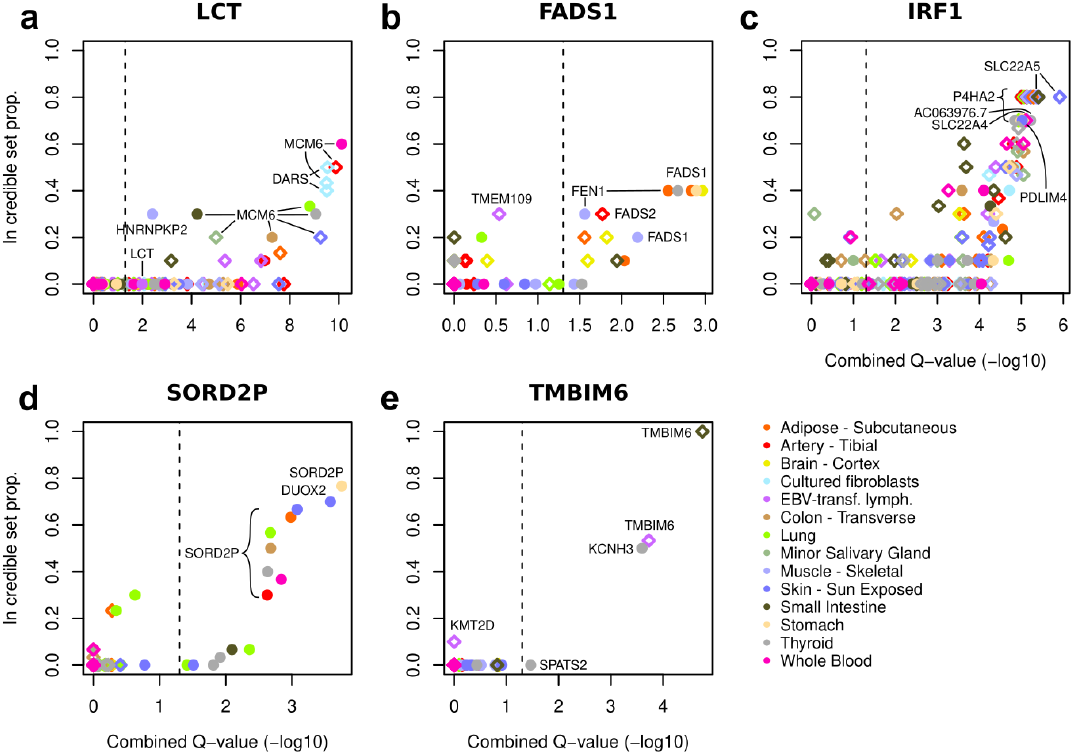
GRe* candidates for selection. **a-e** For five selected loci, the y-axis represents the proportion of 30 SuSiE runs where GRe* features fell in a credible set, while on the x-axis we report the combined Q-values (-log_10_ transformed) across 10 GRe* training runs, the dashed line representing significance cutoff at 5% FDR. Dots represent GReX while diamonds represent GreS, each colored by tissue. As a result there can be multiple diamonds for each gene/tissue representing different splicing features. Labels are present for manually selected genes. Remaining loci are represented in **Figure S7**.

### Transcriptional features in selection-driving credible sets

Among the 10 loci of interest, the *LCT* locus shows the most significant selection signal (**Figure 2A**). The GReX of *MCM6* in whole blood is the feature most consistently appearing in a CS (60% of SuSiE runs) and most significant (Q=7.6e-11), followed by a few splicing features for *MCM6* and *DARS* in artery and cultured fibroblasts. As *LCT* genetic expression and alternative splicing are poorly characterized in GTEx, we could model its expression in only one out of the 14 tissues studied (EBV-transformed lymphocytes). We note, however, that its GReX is among those significantly associated with time (Q=0.01). It is likely that *MCM6* (but also *DARS*) stand out as causal GReX because of their strong anti-correlation with *LCT*. While *MCM6* expression decreases through time, *LCT*’s increases. *The* GReX of *MCM6* also shows the strongest time/space interaction among the candidate genes analyzed (P=1.4e-05 for time/latitude) as its change is markedly stronger in North/North-West Europe (**Figure 3A, S8A**).

**Figure 3:**
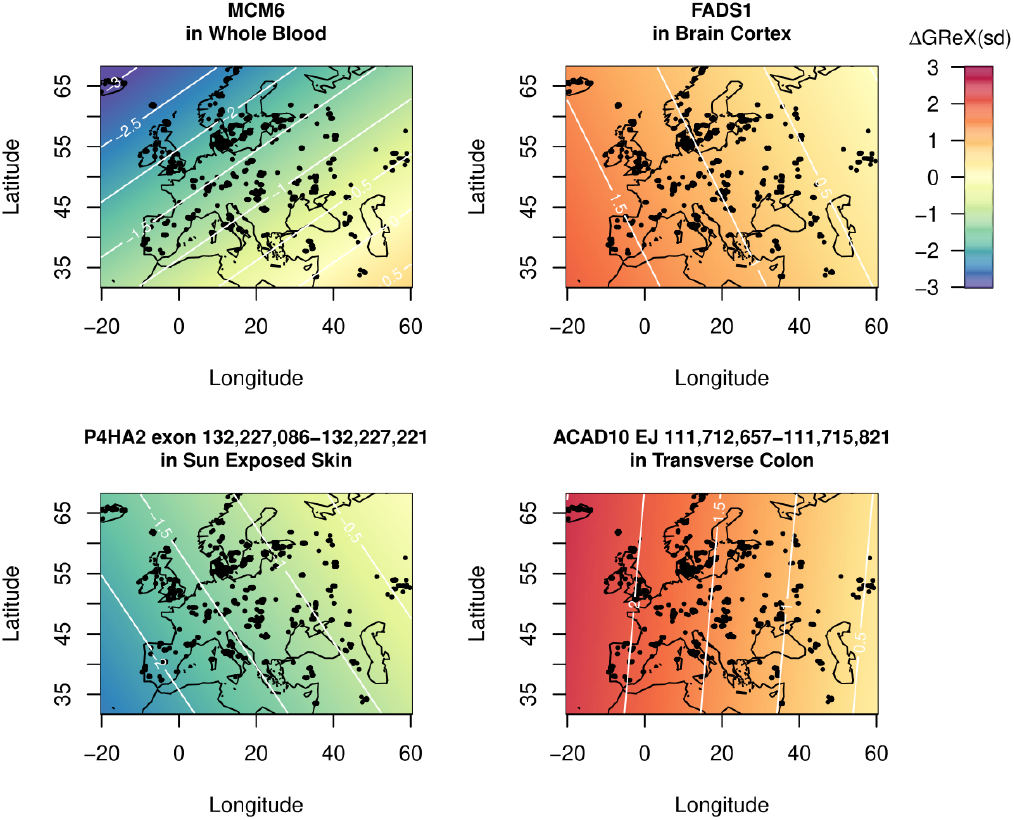
Spatial patterns of GRe* changes. **a-d** Predicted spatial distribution of GRe* change (ΔGRe*) in the last 10.000 years for selected genes. ΔGRe* is reported in units of standard deviation across the discovery sample and is computed with the estimated coefficients for time and its interactions with latitude and longitude (coefficients combined across 10 GRe* training runs). Remaining candidate GRe* are represented in **Figure S8**.

*FADS1*, another gene reported to be under positive selection in several studies^7,14,20^ is also the most consistently appearing in a CS at its locus (**Figure 2B**), although in the initial association analysis the only feature significantly associated with time at this locus was the *TMEM109* GReS (**Figure 1B**). This further underlines the importance of assessing the variability of GRe* models. *FADS1* GReX in four tissues appear in the CS resulting from 40% of SuSiE runs, with Brain Cortex showing the strongest combined significance (Q=1.08e-03). Again, the spatial pattern of selection is uneven, as demonstrated by multiple spatial terms reaching nominal significance, and shown in **Figure 3B**, where we can appreciate a larger increase of *FADS* expression in the west.

We identify a large peak in chr5, previously identified and tagged as *IRF1*^8^ or *PDLIM4*^21^, where the CS inclusion of multiple *SLC22A5* and *P4HA2* GReS in 80% of runs does not allow to precisely finemap the candidate causal transcriptional feature (**Figure 2C**). While a coding variant in *SLC22A4* has been suggested as causal for this selection signal^21^, we observe the insertion of a non-constitutive exon in *P4HA2* 5’UTR (hg38 coordinates chr5:132,227,086-132,227,221), which could lead to an altered translational efficiency of this gene. *P4HA2* is involved in collagen biosynthesis^32^. The insertion of this non-constitutive exon is decreasing over time (in a space-dependent manner, **Figure 3C**) in seven tissues, suggesting that the cause for this selection signal could be multi-factorial.

Another previously documented hit is located at the *OAS3*/*ACAD10* locus. Here a documented adaptive Neanderthal introgression impacted *OAS3*/*OAS1* transcription^33^, but we identify the increased usage of a specific exon junction at *ACAD10* as the most likely adaptive candidate (**Figure S7A**, especially in the west, **Figure 3D**). This is consistent with the finding by Irving-Pease et al^8^ that a variant within its 13th intron is the one with the highest selection signal at this locus in the same ancient dataset.

All novel hits except for *DNAJB12* show large concordance across runs, with the genes more often appearing in CS doing so at least 77% of times. A decreased expression of *SORD2P* GReX in the stomach emerged as the most likely selected feature at its locus (**Figure 2D**).

*SORD2P* is a noncoding pseudogene paralogous to *SORD*, located less than 140 kb downstream. Although noncoding, its transcript may exert a regulatory effect on *SORD*, which converts sorbitol to fructose^34^ and has been implicated in Charcot-Marie-Tooth disease^35^. *AC012370*.*3*, together with its antisense *AC074391*.*1* are two noncoding transcripts whose expression decreases over time in thyroid are both the most likely selected features at their locus (**Figure S7B**), although their function is yet uncertain. Finally, *TMBIM6* alternative splicing alteration in the small intestine is the only feature falling in a CS in 100% of runs (**Figure 2E**): this gene encodes for an endoplasmic-reticulum protein that regulates calcium homeostasis and suppresses apoptosis during stress^36^. Variants at this gene have been associated with bone mineral density^37^.

## Discussion

We applied expression and splicing predictive models to samples covering the last ten thousand years across Western Eurasia, and by modeling the predicted molecular traits with time and space we identified and refined a total of 11 signals of selection acting on transcriptional regulation. Expanding both sample set and probed transcriptional features with respect to a previous analysis in Britain^21^, we were able to recognize two more known selected loci as transcription-mediated (*TLR* and *GATA4*) and identify four novel loci, although we could not replicate their signals at *SLC44A5* and *NUP85*. We did not find evidence of a role of expression or splicing for 11 remaining known selection candidates, either because the adaptation does not go through regulatory mechanisms or because it takes place in a very specific cellular context.

While the study referenced above did not include genetic and spatial covariates, the wide-ranging dataset we analyzed required their introduction. We chose a conservative approach but it is likely that modeling population structure with less than 10 principal components or as a random effect (as done in a recent study^25^) could lead to a higher sensitivity, at the cost of finding signals potentially explained by genetic drift or incoming migration. While precisely disentangling genetic changes due to selection or demography remains a challenge, with an extensive modeling of the dataset heterogeneity we were still able to capture robust transcriptional changes and to spatially characterize those previously known, such as *MCM6, FADS1, ACAD10* and *P4AH2*.

By exploiting an imputed genotype dataset we could apply full predictive models as it’s done on contemporary samples, which allows a moderate but significant predictivity improvement over downsampled models previously built for ancient data^20^. This gain holds when validating on genetically similar (European) samples and on non-European samples, which is crucial given millennia of migrations and drift between modern and ancient Europeans. Including alternative splicing as an orthogonal transcriptional dimension, while having the potential of pinpointing more precisely the transcriptional feature under selection, did not reveal radically independent signals and remains more complex to interpret.

The loci discovered spanned multiple genes and contexts, tied by the linkage disequilibrium between the respective predictive variants. For the first time to our knowledge we attempted to resolve these signals of selection by applying fine-mapping technologies derived from the GWAS field. This was coupled with a repeated re-training of the GRe* predictive models to account for their uncertainty, which revealed a sizable variability across associations resulting from different GRe* training runs. The application of methods able to inherently model uncertainty in genetic variables (e.g. tissue-gene fine-mapping^31^), and connecting them with multiple, interacting, non-genetic factors, can improve our approach and pave a new way to identify adaptation-driving functional features.

The difference in predictability of transcriptional features across tissues and genes represents a challenge for our methodology. Many genes were excluded altogether from the analysis due to a poorly predictive GRe* model. Nevertheless, the correlation of each GRe* with the corresponding transcriptional trait depends not only on the prediction error, which is a function of training sample size and explains inter-tissue differences at the same gene; but also on its heritability. Heritability also defines the potential response of a trait to selection, which is larger for highly heritable traits. This suggests that well-predicted GRe* values, at least for recent, unfixed adaptative features, are also more likely to be the real drivers of natural selection. Molecular trait imputation could therefore naturally highlight features with higher potential of being the selection targets, provided that the correct cellular context is included.

In conclusion, the analysis of genetically regulated transcriptional phenotypes combined with a spatio-temporally resolved dataset and fine-mapping allowed us to gain mechanistic insight into candidate loci for selection. The training of genetic models for additional molecular phenotypes and biological contexts, unlocked by the deluge of molecular data such as protein abundance and cell-type resolution, implies that the potential of this approach will continue to grow.

## Methods

### Building molecular traits prediction models

We developed an in-house implementation of PrediXcan^16^ to build prediction models of GReX and GReS trained on GTEx v8 data^13^. We used whole-genome VCFs mapped on GRCh38 and expression or intron excision ratios made available by the consortium. We selected 14 tissues of interest maximizing sample size and tissue diversity. Variant and sample selection, inclusion of covariates and model training were performed matching the methods in prediXcan^16^ with the only exception of the hyper-parameters being tuned using threefold cross-validation. We trained our models using a randomly selected 80% subset of European samples from the GTEx dataset. The remaining 20% subset was used to evaluate prediction accuracy, measured as Pearson correlation coefficient (R) between predicted and actual GRe* values in the test set. Accuracy was also measured in non-European samples.

To compare our results to previous work^20^ we also generated prediction models for the Skin Sun Exposed Lower leg tissue using only the sites from the GTEx dataset in the 1240k variants assayed in ancient humans from the AADR (V54.1.p1, Dataverse 8.0)^22^. Once we filtered out the GTEX vcf for those sites, we followed the same approach to generate the models as specified above.

### Prediction of molecular traits on the ancestral genomes

We predicted GReX and GReS on imputed, non-flagged ancient genomes from western Eurasia (ancient dataset, n=1,145)^6^. We also predicted these molecular traits on 80 high-coverage modern European genomes (20 each from FIN, GBR, IBS, TSI to maintain balance with ancient sample groups) from the 1000 Genomes project^24^ (**Table S1, Figure S1**). We predicted molecular traits on the intersection of SNPs from these two datasets (discovery set, n. SNPs=25,326,340) for each GRe* feature with prediction models having R greater than 0.1 and gene-level variant imputation quality score (computed as in Poyraz et al.^21^) greater than 0.8.

### Regressing on spatio-temporal axes

We modelled how the predicted molecular traits for the ancient samples vary depending on a set of predictors. In addition to the 10 first PCs computed for the discovery dataset, we included latitude, longitude, and time as predictors as well as latitude:longitude, latitude:time and longitude:time interaction terms. Time is defined as Age Average in **Table S1**. PCs were computed with the “bigsnipr” package following the best practices described in Prive’ et al^38^. For each gene for which we had the prediction in at least one tissue, we aggregated the P-value for each spatio-temporal predictor across tissues using Aggregated Cauchy Association Test (ACAT)^26^. Gene significance was assessed at FDR=0.05 independently for GReX and GReS.

### Candidate loci for selection

We harvested candidate loci for selection from six publications^7,8,15,21,27,28^ performing selection scans in the period and geographic space considered in our work, lifted to GRCh38 and enlarged them 1Mb per side. Eighteen loci found overlapping across at least two of these publications were merged and considered as replicable candidates of selection, see **Table S5**. New hits identified in our analysis—defined as regions within 1 Mb upstream and downstream of significant GRe*/time associations—were added to the list.

### Loci refinement and fine-mapping

GRe* models passing the filters on R and imputation quality score described in the previous sections were re-trained 10 times, each time leaving out a random 20% of the GTEx training set, in order to assess variability across models.

P-values across runs were combined with the goal to emphasize consistency while accounting for the average 64% training set overlap and the usage of the same discovery set in all runs. We therefore selected the average Z score as a conservative approach. Note that a classic Stouffer’s Z meta-analysis approach would reduce to that when considering all equal weights for individual analyses and full sample overlap. To estimate ΔGRe* we employed the combined coefficients for time, latitude, longitude and their interactions and sampled a matrix of coordinates across Western Eurasia at 10 thousand years distance, then computing the difference in fitted expression.

For the fine-mapping we used summary-based SuSiE^30^, with the associations with time of each GRe* as summary statistics. LD matrices were computed as correlation matrices between GRe* in the discovery dataset. We ran SuSiE separately for each of the ten different GRe* resampling iterations, and iterating the L parameter (maximum allowed number of independent signals) from 1 to 3, resulting in 30 independent runs across which we used as read-out the fraction of runs in which each GRe* was included in a CS.

## Data availability

GTEx v8 gene expression and alternative splicing normalised measurements were accessed from the project portal at https://www.gtexportal.org/home/downloads/adult-gtex/qtl. Individual-level data for GTEx subjects were obtained from dbGAP (https://www.ncbi.nlm.nih.gov/gap/) under project #18670. GENCODE annotation release 26 for GRCh38 was obtained from https://www.gencodegenes.org/human/release_26.html. We accessed the imputed genotypes for no flag ancient individuals used in this study at https://doi.org/10.17894/ucph.d71a6a5a-8107-4fd9-9440-bdafdfe81455. The high-coverage genomes from the 1000 Genomes project were downloaded from the ftp site linked at https://www.internationalgenome.org/data-portal/data-collection/30x-grch38.

## Acknowledgements

This work was supported in part by a Fondazione CRT grant (grant 2021.1787 to P.P.). We thank Renato Polimanti, Alessandro Raveane and Daniela Fusco for insightful discussions.

## Author contributions

D.M. and P.P. conceived and supervised the project; M.A. and D.M. performed the data analysis; M.A. and D.M. wrote the manuscript with the contribution of all the co-authors.

## References

1. Lazaridis, I. et al. Ancient human genomes suggest three ancestral populations for present-day Europeans. Nature 513, 409–13 (2014).

2. Lipson, M. et al. Parallel palaeogenomic transects reveal complex genetic history of early European farmers. Nature 551, 368–372 (2017).

3. Posth, C. et al. Palaeogenomics of Upper Palaeolithic to Neolithic European hunter-gatherers. Nature 615, 117–126 (2023).

4. Speidel, L. et al. High-resolution genomic history of early medieval Europe. Nature 637, 118–126 (2025).

5. Haak, W. et al. Massive migration from the steppe was a source for Indo-European languages in Europe. Nature 522, 207–211 (2015).

6. Allentoft, M. E. et al. Population genomics of post-glacial western Eurasia. Nature 625, 301–311 (2024).

7. Mathieson, I. et al. Genome-wide patterns of selection in 230 ancient Eurasians. Nature 528, 499–503 (2015).

8. Irving-Pease, E. K. et al. The selection landscape and genetic legacy of ancient Eurasians. Nature 625, 312 (2024).

9. Marnetto, D. et al. Ancestral genomic contributions to complex traits in contemporary Europeans. Curr. Biol. 32, 1412-1419.e3 (2022).

10. Pankratov, V. et al. Ancestral genetic components are consistently associated with the complex trait landscape in European biobanks. Eur. J. Hum. Genet. https://doi.org/10.1038/s41431-024-01678-9 (2024) doi:10.1038/s41431-024-01678-9.

11. Grossman, S. R. et al. Identifying recent adaptations in large-scale genomic data. Cell 152, 703–713 (2013).

12. Aguet, F. et al. Molecular quantitative trait loci. Nat. Rev. Methods Primer 3, 4 (2023).

13. THE GTEX CONSORTIUM. The GTEx Consortium atlas of genetic regulatory effects across human tissues. Science 369, 1318–1330 (2020).

14. Buckley, M. T. et al. Selection in Europeans on Fatty Acid Desaturases Associated with Dietary Changes. Mol. Biol. Evol. 34, 1307–1318 (2017).

15. Kerner, G. et al. Genetic adaptation to pathogens and increased risk of inflammatory disorders in post-Neolithic Europe. Cell Genomics 3, 100248 (2023).

16. Gamazon, E. R. et al. A gene-based association method for mapping traits using reference transcriptome data. Nat. Genet. 47, 1091–1098 (2015).

17. Barbeira, A. N. et al. Exploring the phenotypic consequences of tissue specific gene expression variation inferred from GWAS summary statistics. Nat. Commun. 9, 1825 (2018).

18. Zhou, D. et al. A unified framework for joint-tissue transcriptome-wide association and Mendelian randomization analysis. Nat. Genet. 52, 1239–1246 (2020).

19. Ferrando-Bernal, M., Brand, C. M. & Capra, J. A. Inferring human phenotypes using ancient DNA: from molecules to populations. Curr. Opin. Genet. Dev. 90, 102283 (2025).

20. Colbran, L. L., Johnson, M. R., Mathieson, I. & Capra, J. A. Tracing the Evolution of Human Gene Regulation and Its Association with Shifts in Environment. Genome Biol. Evol. 13, evab237 (2021).

21. Poyraz, L., Colbran, L. L. & Mathieson, I. Predicting Functional Consequences of Recent Natural Selection in Britain. Mol. Biol. Evol. 41, msae053 (2024).

22. Mallick, S. et al. The Allen Ancient DNA Resource (AADR) a curated compendium of ancient human genomes. Sci. Data 11, 182 (2024).

23. The 1000 Genomes Project Consortium et al. A global reference for human genetic variation. Nature 526, 68–74 (2015).

24. Byrska-Bishop, M. et al. High-coverage whole-genome sequencing of the expanded 1000 Genomes Project cohort including 602 trios. Cell 185, 3426-3440.e19 (2022).

25. Akbari, A. et al. Pervasive findings of directional selection realize the promise of ancient DNA to elucidate human adaptation. Preprint at 10.1101/2024.09.14.613021 (2024).

26. Liu, Y. et al. ACAT: A Fast and Powerful p Value Combination Method for Rare-Variant Analysis in Sequencing Studies. Am. J. Hum. Genet. 104, 410–421 (2019).

27. Field, Y. et al. Detection of human adaptation during the past 2000 years. Science 354, 760–764 (2016).

28. Palamara, P. F., Terhorst, J., Song, Y. S. & Price, A. L. High-throughput inference of pairwise coalescence times identifies signals of selection and enriched disease heritability. Nat. Genet. 50, 1311–1317 (2018).

29. Sabeti, P. C. et al. Genome-wide detection and characterization of positive selection in human populations. Nature 449, 913–8 (2007).

30. Zou, Y., Carbonetto, P., Wang, G. & Stephens, M. Fine-mapping from summary data with the “Sum of Single Effects” model. PLOS Genet. 18, e1010299 (2022).

31. Strober, B. J., Zhang, M. J., Amariuta, T., Rossen, J. & Price, A. L. Fine-mapping causal tissues and genes at disease-associated loci. Nat. Genet. 57, 42–52 (2025).

32. Myllyharju, J. Prolyl 4-hydroxylases, the key enzymes of collagen biosynthesis. Matrix Biol. 22, 15–24 (2003).

33. Sams, A. J. et al. Adaptively introgressed Neandertal haplotype at the OAS locus functionally impacts innate immune responses in humans. Genome Biol. 17, 246 (2016).

34. El-Kabbani, O., Darmanin, C. & Chung, R. Sorbitol Dehydrogenase: Structure, Function and Ligand Design. Curr. Med. Chem. 11, 465–476 (2004).

35. Cortese, A. et al. Biallelic mutations in SORD cause a common and potentially treatable hereditary neuropathy with implications for diabetes. Nat. Genet. 52, 473–481 (2020).

36. Lebeaupin, C., Blanc, M., Vallée, D., Keller, H. & Bailly-Maitre, B. BAX inhibitor-1: between stress and survival. FEBS J. 287, 1722–1736 (2020).

37. He, D. et al. A longitudinal genome-wide association study of bone mineral density mean and variability in the UK Biobank. Osteoporos. Int. 34, 1907–1916 (2023).

38. Privé, F., Luu, K., Blum, M. G. B., McGrath, J. J. & Vilhjálmsson, B. J. Efficient toolkit implementing best practices for principal component analysis of population genetic data. Bioinformatics 36, 4449–4457 (2020).

