## Supplementary Figures for "Imputed transcriptional patterns in ancient genomes reveal molecular targets of selection"

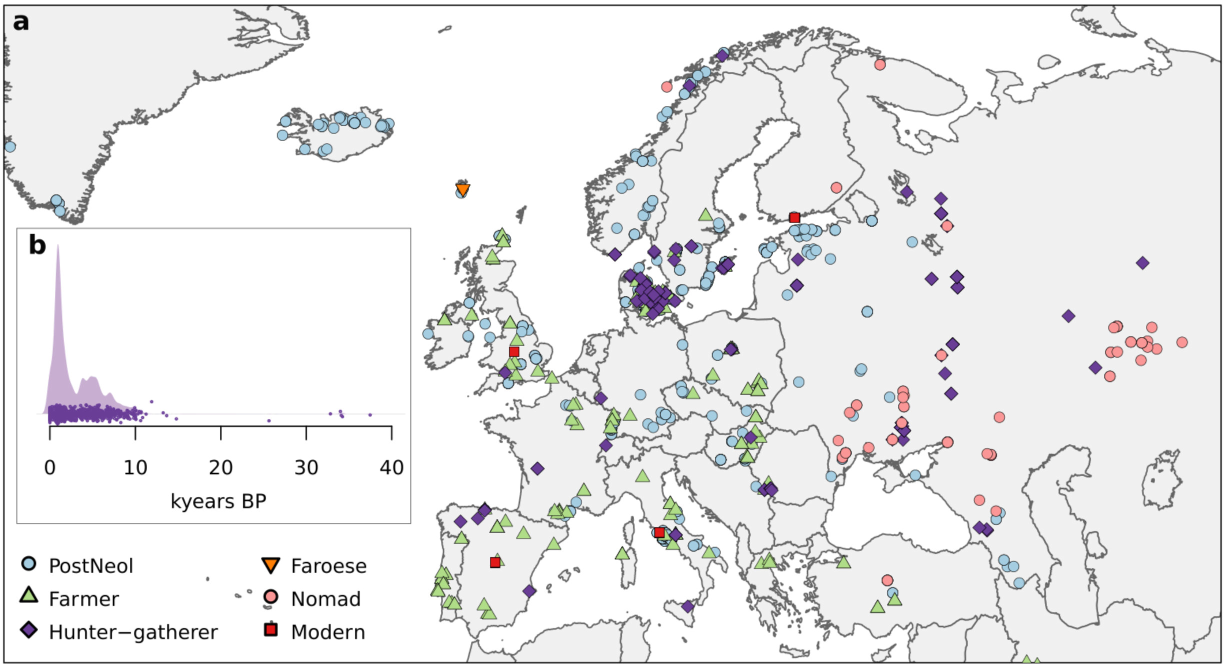


***Figure S1. Ancient Samples. a.*** *Map of the sampling places of the discovery dataset.* ***b.*** *Time distribution of the samples in the discovery dataset*

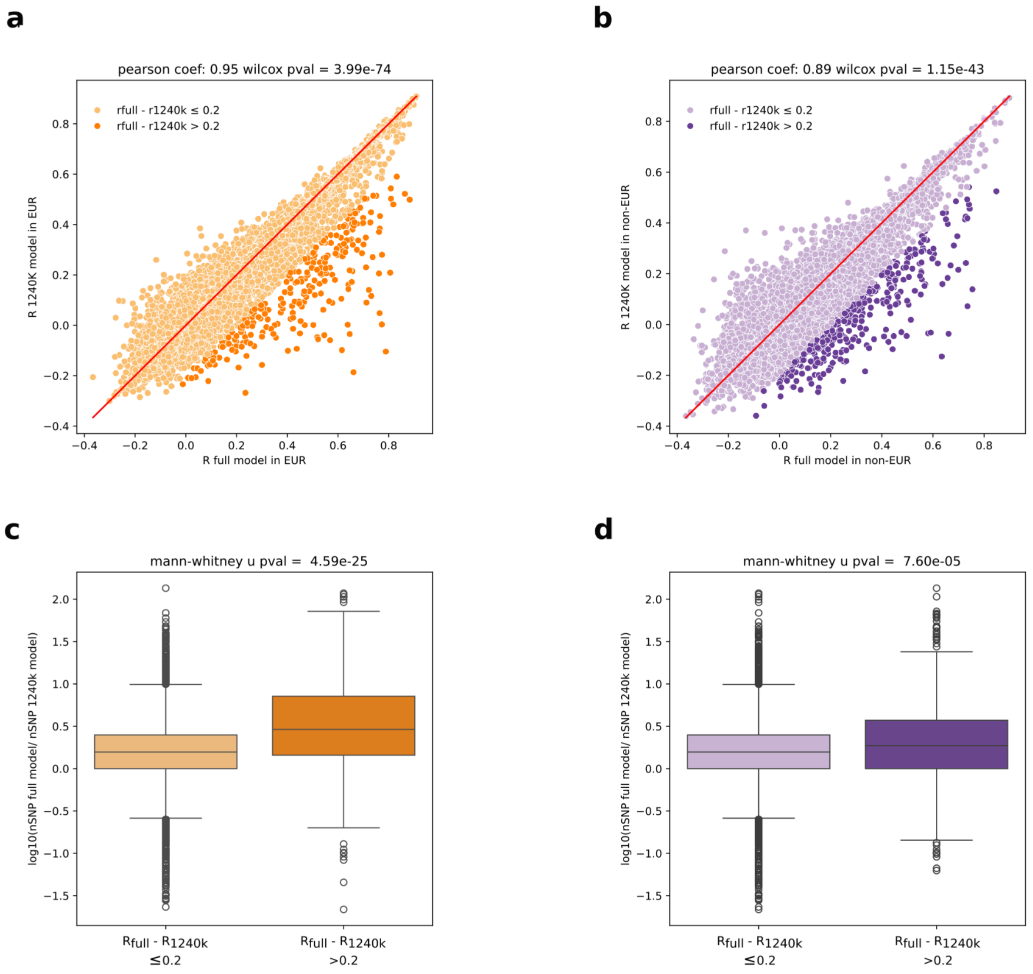


***Figure S2. Comparison between full vs 1240k downsampled model***. ***a.*** *Correlation of Skin Sun Exposed Lower leg GReX prediction performance (R) in GTEx European samples using models trained on the full variant set versus the 1240k downsampled variant set.* ***b.*** *Correlation of Skin Sun Exposed Lower leg GReX prediction performance (R) in GTEx non-European samples using models trained on the full variant set versus the 1240k downsampled variant set.* ***c.*** *Proportion of SNPs present in the full GTEx training set but absent in the 1240k training set, for genes where prediction R is ≥20% higher in the full Skin Sun Exposed Lower leg model compared to the 1240k model in European individuals.* ***d.*** *Same as* ***(c)****, but for predictions in non-European individuals.*

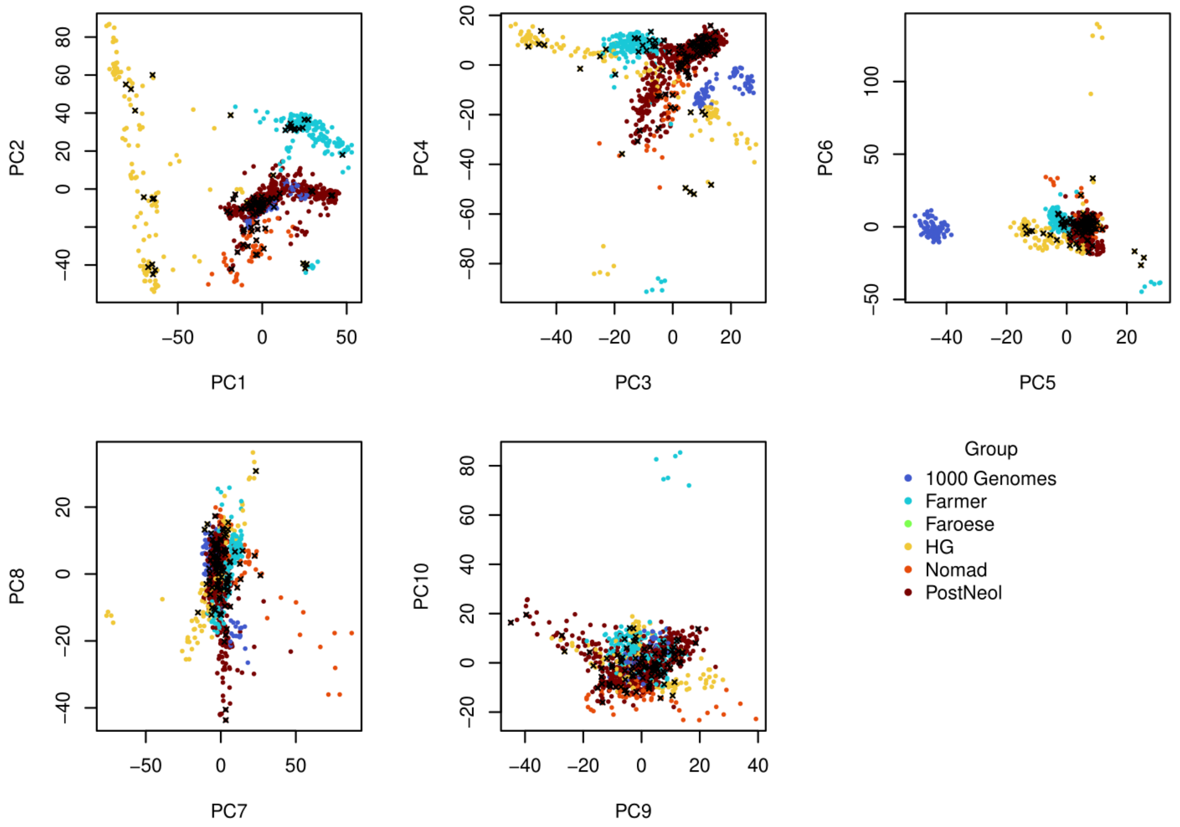


***Figure S3. Discovery Set Genetic Principal Components.*** *Each ancient sample is colored by lifestyle group as defined in the original data, with 1000 Genomes samples added as an extra group. Black crosses indicate outlier or related samples not used in the PC calculation.*

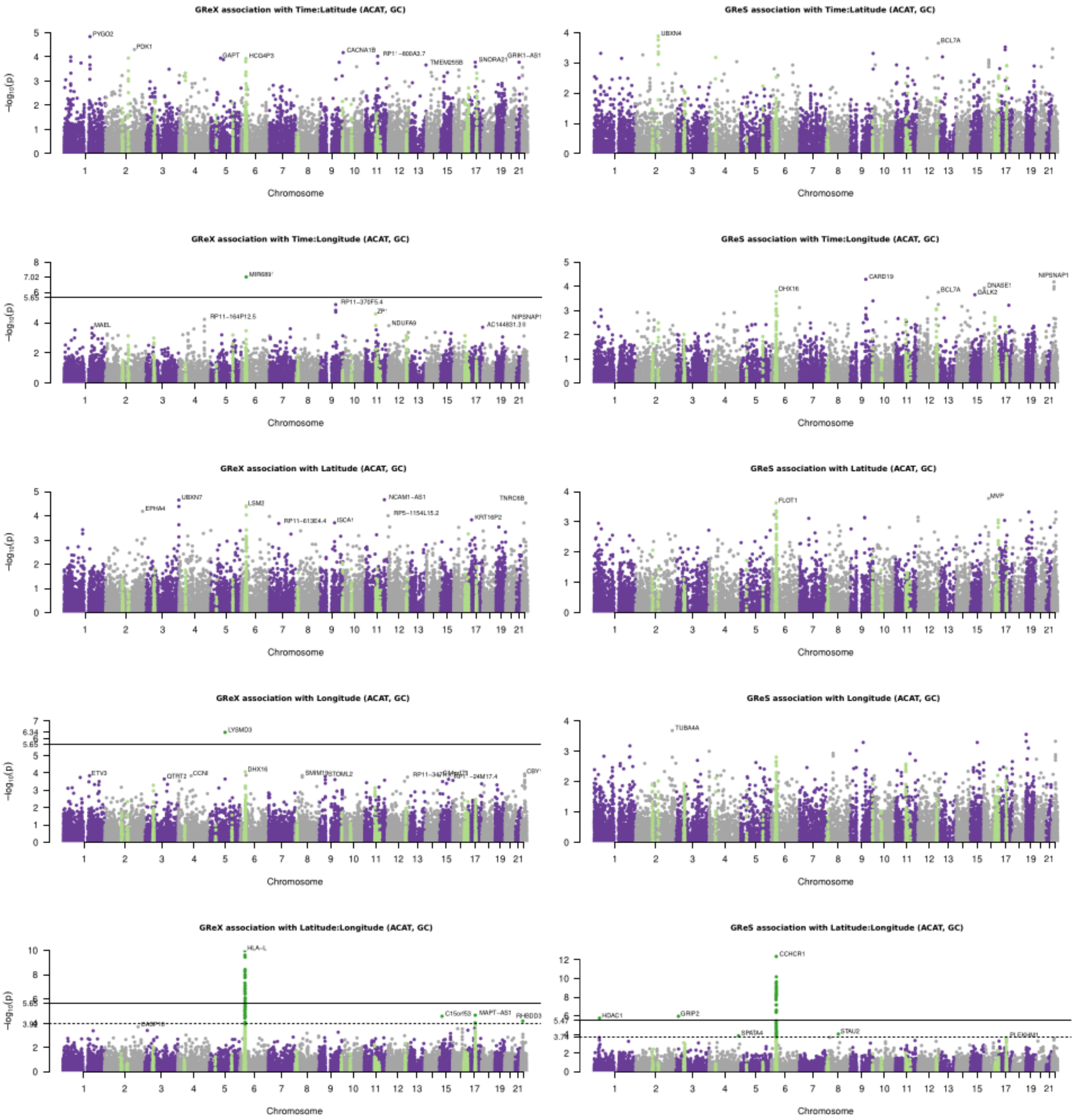


***Figure S4*. Manhattan plots for Latitude, Longitude, and their interactions with time.** *P-values represented are treated as in* ***Figure 1A*** *and* ***B****, that is, resulting from ACAT aggregation across tissues and after transformation with Genomic Control. Solid and dashed lines, when present, identify significance after Bonferroni correction and at 5% FDR, respectively.*

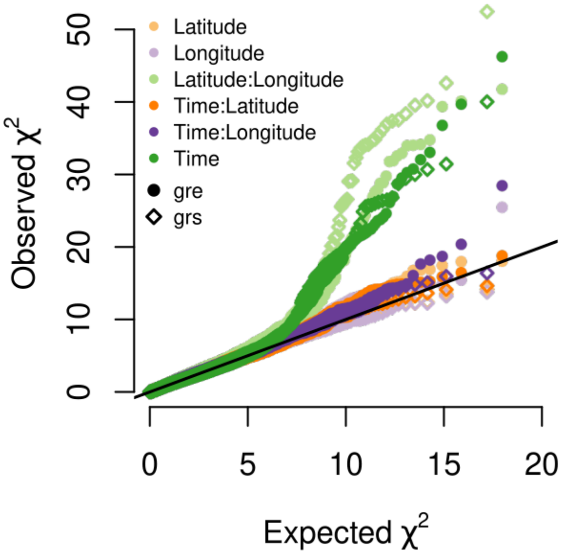


**Figure S5**. *QQ plot of $X^2$ for each predictor separately for GReX (dots) and GReS (diamonds). Same as **Fig 2C** but without removing the HLA region.*

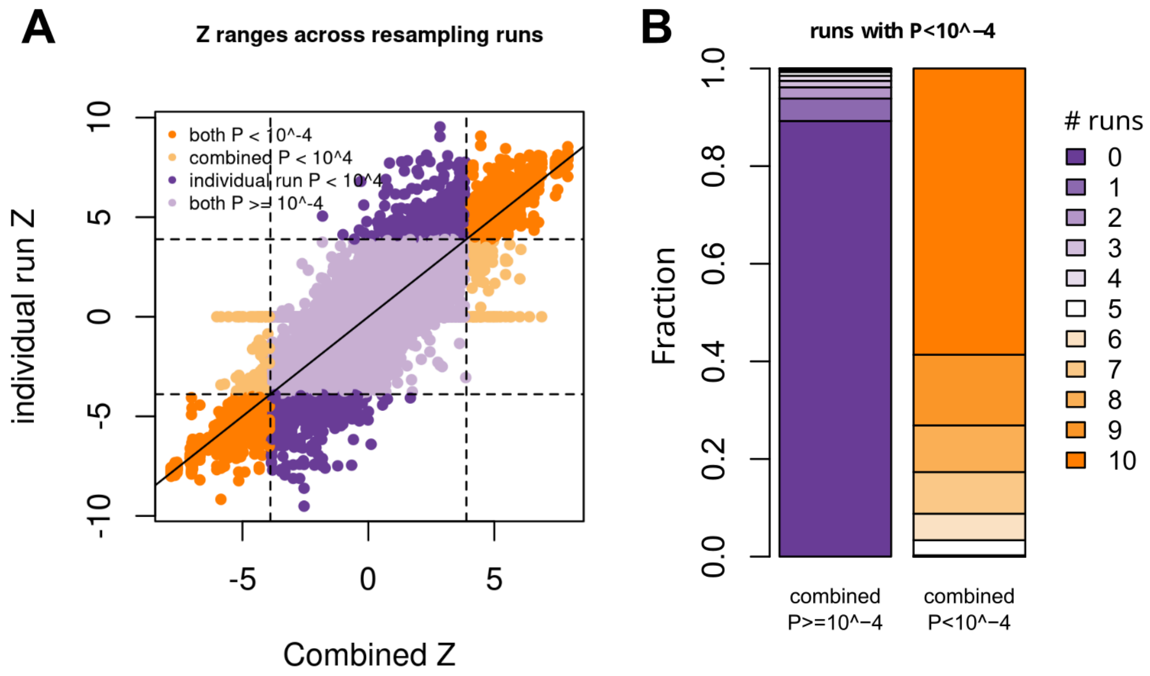


***Figure S6. a.*** *Each dot represents an individual training run for each GRe* among those falling within the 10 candidate loci re-trained with resampling, placed along the Y axis according to its Z for association with time. On the X axis is represented the combined Z across resampling runs for the same coefficient. Each dot is colored with respect to the significance of the individual run it represents and of the combined Z score, according to a representative threshold of* $P=10^{-4}$ *(double-sided), corresponding to an absolute Z of =3.89, indicated by the dashed lines.* ***b.*** *Proportion of GRe*, separating for significant and not significant combined Z, that reached significance in # independent GRe* training runs. 5848 and 387 GRe* are represented in the right and left column respectively, for a total of 6235 re-trained features. Significance is assessed according to the same threshold as the previous panel.*

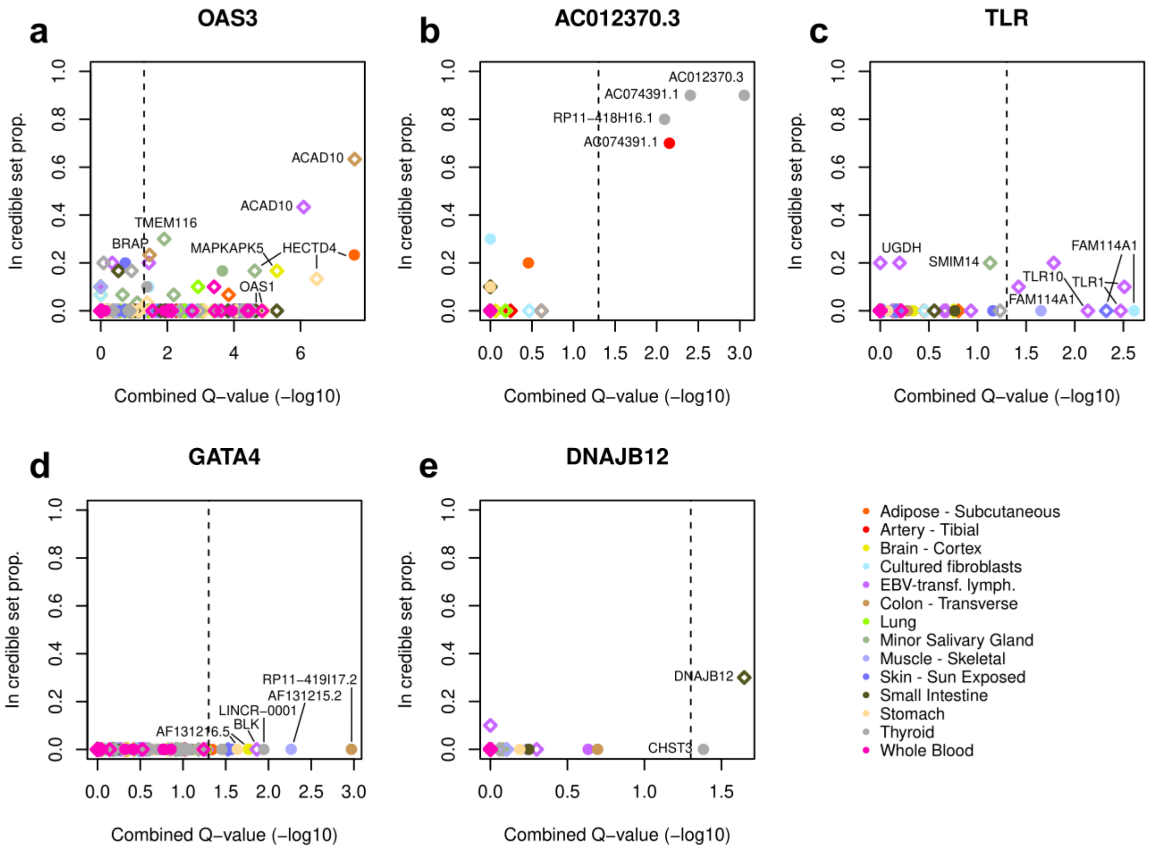


***Figure S7. Other GRe* candidates for selection.*** *For loci not represented in **Figure 2**, the y-axis represents the proportion of 30 SuSiE runs where GRe* features fell in a credible set, while on the x-axis we report the combined Q-values (-log$_{10}$ transformed) across 10 GRe* training runs, the dashed line representing significance cutoff at 5% FDR. Dots represent GReX while diamonds represent GreS, each colored by tissue. As a result there can be multiple diamonds for each gene/tissue representing different splicing features. Labels are present for manually selected genes.*

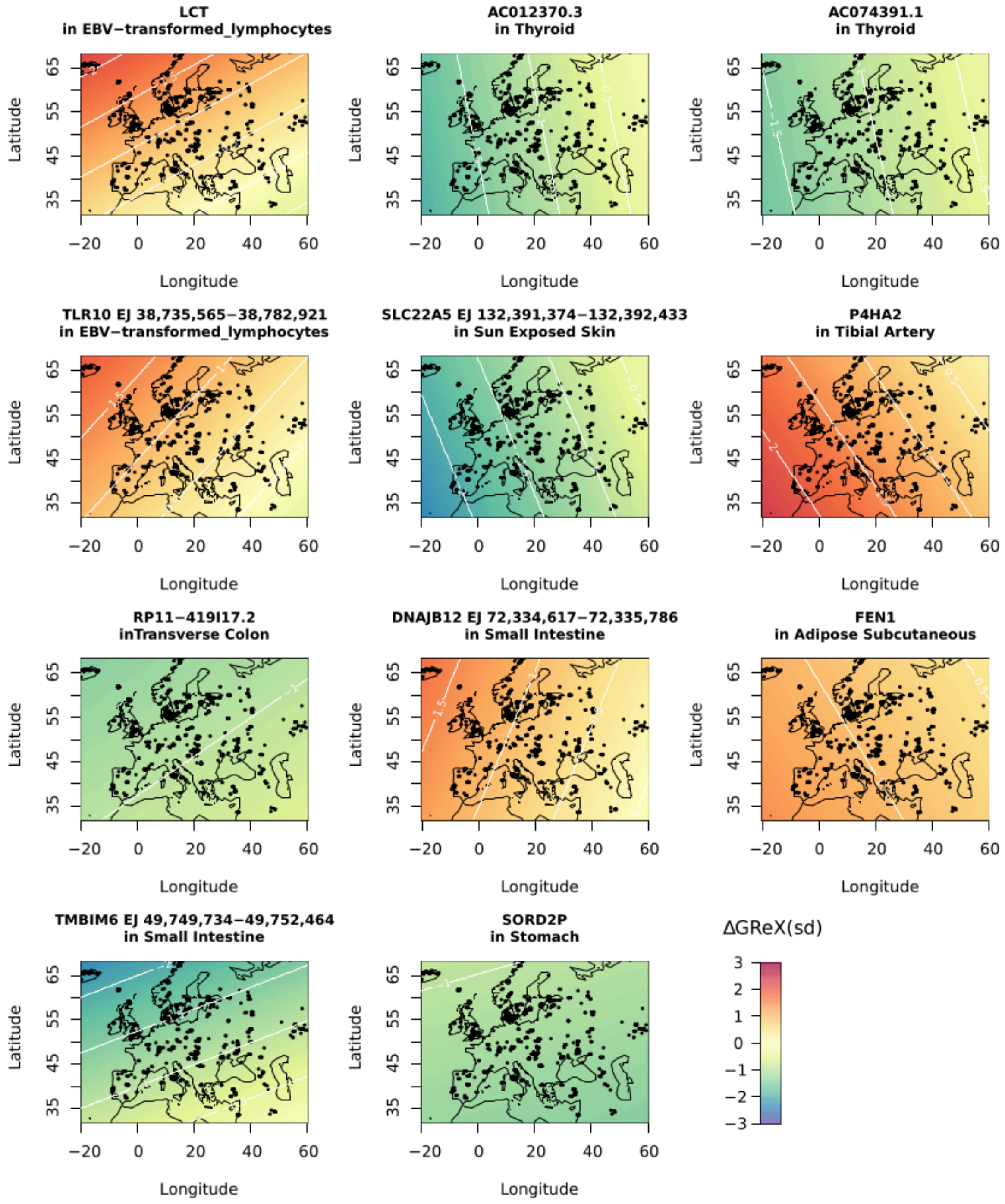


***Figure S8: Other spatial patterns of GRe* changes.*** *Predicted spatial distribution of GRe* change (ΔGRe*) in the last 10.000 years for genes not represented in* ***Figure 3****. ΔGRe* is reported in units of standard deviation across the discovery sample and is computed with the estimated coefficients for time and its interactions with latitude and longitude (coefficients combined across 10 GRe* training runs).*
